# SARS-CoV-2 infection induces a pro-inflammatory cytokine response through cGAS-STING and NF-κB

**DOI:** 10.1101/2020.07.21.212639

**Authors:** Christopher J. Neufeldt, Berati Cerikan, Mirko Cortese, Jamie Frankish, Ji-Young Lee, Agnieszka Plociennikowska, Florian Heigwer, Sebastian Joecks, Sandy S. Burkart, David Y. Zander, Mathieu Gendarme, Bachir El Debs, Niels Halama, Uta Merle, Michael Boutros, Marco Binder, Ralf Bartenschlager

## Abstract

SARS-CoV-2 is a novel virus that has rapidly spread, causing a global pandemic. In the majority of infected patients, SARS-CoV-2 leads to mild disease; however, in a significant proportion of infections, individuals develop severe symptoms that can lead to permanent lung damage or death. These severe cases are often associated with high levels of pro-inflammatory cytokines and low antiviral responses which can lead to systemic complications. We have evaluated transcriptional and cytokine secretion profiles from infected cell cultures and detected a distinct upregulation of inflammatory cytokines that parallels samples taken from infected patients. Building on these observations, we found a specific activation of NF-κB and a block of IRF3 nuclear translocation in SARS-CoV-2 infected cells. This NF-κB response is mediated by cGAS-STING activation and could be attenuated through STING targeting drugs. Our results show that SARS-CoV-2 curates a cGAS-STING mediated NF-κB driven inflammatory immune response in epithelial cells that likely contributes to inflammatory responses seen in patients and might be a target to suppress severe disease symptoms.

## Introduction

In late 2019 SARS-CoV-2 emerged as a highly infectious coronavirus that causes respiratory disease in humans, termed COVID-19. Since the initial identification, SARS-CoV-2 has spread around the world leading the World Health Organization to declare a pandemic. SARS-CoV-2 infection causes respiratory symptoms that range from mild to severe and can result in lasting lung damage or death in a significant number of cases ^1^. One of the hallmarks of severe COVID-19 disease is low levels of type I interferons (IFNs) and overproduction of inflammatory cytokines such as IL-6 and TNF ^2–5^. This unbalanced immune response fails to limit virus spread and can cause severe systemic symptoms ^3,6^. Therapies aimed at modulating immune activation to attenuate the detrimental inflammatory response or promote an antiviral cytokine response represents an important avenue for treating patients with severe COVID-19.

SARS-CoV-2 is a plus-strand RNA virus that replicates its genome in the cytosolic compartment of the cells. Like all plus-strand RNA viruses, this replication process requires the production of a negative-strand RNA template in order to amplify the positive sense viral genome. This process and probably also the production of subgenomic RNAs of negative and positive polarity, produces double strand (ds)RNAs that can be sensed by cytosolic immune receptors (pattern recognition receptors, PRRs) that subsequently activate antiviral pathways ^7^. In addition to direct viral sensing, cells have also evolved ways to detect the indirect effects of virus infection, such as nuclear or mitochondrial damage caused by the heavy cellular burden of virus replication. Cytoplasmic DNA sensors including cGAS-STING, IFI16, or AIM2, recognize dsDNA from DNA viruses, but have also been shown to play an important role in RNA virus infection, either through directly recognising viral signatures or through sensing of cellular DNA released from mitochondria or nuclei due to cellular stress (reviewed in ^8,9^). Substrate recognition by either RNA or DNA sensors leads to signalling cascades that activate two major branches of the innate immune response, the type I/III IFN response and the inflammatory cytokine response. The type I/III IFN pathways are directly involved in protecting neighboring cells from virus spread and are vital for the immediate cell-intrinsic antiviral response. The inflammatory cytokine response is involved in recruitment and activation of immune cells, which is required to initiate an adaptive immune response.

Due to the effective nature of innate immune sensing and responses plus-strand RNA viruses have evolved numerous ways to limit or block these cellular pathways. For many viruses, the initial line of defence is to hide viral replication intermediates within membrane compartments that block access to cytosolic PRRs, such as RIG-I or MDA5 ^10^. In the case of coronaviruses, this is achieved through the formation of replication organelles composed predominantly of double membrane vesicles, within which viral RNA replication occurs ^11,12^. Coronaviruses can also evade recognition by immune receptors through modification of viral RNA to resemble host mRNA^13–16^. In addition to these passive immune evasion strategies, coronaviruses utilize various mechanisms to actively target and block key immune sensors or signalling molecules (reviewed in ^17,18^). For SARS-CoV-1, a closely related virus, several viral proteins have been shown to block RIG-I/MDA5 sensing, as well as the downstream activation of TBK1 and IRF3 ^19–24^. SARS-CoV-1 also efficiently blocks IFN receptor and JAK-STAT signalling to stop downstream immune activation ^19,25–27^. Additionally, the SARS-CoV-1 papain-like protease (PLP) has been shown to interfere with cGAS-STING activation also limiting activation of innate immune pathways ^28^. The combination of these actions can lead to an imbalance between proinflammatory and antiviral immune responses.

Although numerous immune evasion mechanisms have been characterized for other pathogenic coronaviruses, it remains to be determined whether similar processes exist for SARS-CoV-2. Given the homology of SARS-CoV-1 to SARS-CoV-2, they may have many conserved antagonistic strategies, however, key differences in infection and disease could suggest divergent pathways. Early reports on SARS-CoV-2 demonstrated that infection is highly sensitive to type I/III IFN treatment ^29–32^. In combination with the low levels of IFN reported to be secreted in severe cases, this suggests that like SARS-CoV-1, SARS-CoV-2 infection actively blocks immune activation. Transcriptomic analyses of SARS-CoV-2 infected cells generated ambiguous results on the induction of type I/III IFNs and the subsequent expression of IFN stimulated genes (ISGs). On the one hand, it was shown that SARS-CoV-2 triggers only an attenuated immune response suggesting a block in PRR signaling pathways, which would parallel SARS-CoV-1 and MERS-CoV infections ^29^. On the other hand, several studies argue for a strong induction of IFN responses in both lung and intestinal infection models ^30,33^. Additionally, proteomics approaches determining SARS-CoV-2 protein interactions with host factors in exogenous expression conditions revealed several interactions with key immune regulators including MAVS, TBK1 and several co-factors involved in IRF3 activation ^34,35^. However, many of these findings are still observational leaving the mechanisms of SARS-CoV-2 innate immune response modulation unresolved.

Here, we report the transcriptomic profiles derived from SARS-CoV-2 infected human lung cells showing a specific bias towards an NF-κB mediated inflammatory response and a restriction in the TBK1 specific IRF3/7 activation and subsequent IFN response. Consistently, secreted cytokine profiles from both severe COVID-19 patients and SARS-CoV-2 infected lung epithelial cells, were enriched for pro-inflammatory cytokines and lacked type I/III IFNs. We also demonstrate that SARS-CoV-2 infection leads specifically to NF-κB but not IRF3 nuclear localization and that poly(I:C)-induced pathway activation is attenuated in infected cells. Finally, we show that the cGAS-STING pathway is activated by SARS-CoV-2 infection, leading to a specific NF-κB response and that inflammatory cytokine upregulation can be mitigated by STING inhibitory drugs. These results provide insight into how innate immune responses are modulated by SARS-CoV-2 in epithelial cells likely contributing to the strong inflammatory responses observed in severe COVID-19 cases.

## Results

### Kinetics of SARS-COV-2 infection in lung epithelial cells

SARS-CoV-2 predominantly infects airway and lung tissue in infected individuals. In order to determine the effects of SARS-CoV-2 on human lung epithelial cells, Calu-3 and A549 cells were infected with SARS-CoV-2 and virus growth, as well as host transcriptional architecture was determined over a time course of infection. In contrast to Calu-3, A549 cells lack endogenous expression of the major SARS-CoV-2 entry receptor ACE2 and, hence, are not naturally permissive to SARS-CoV-2 infection ^36^. We therefore used an engineered A549 cell line stably expressing ACE2 (A549-ACE2), which is susceptible to SARS-CoV-2 infection ^12,37^. For both, A549-ACE2 and Calu-3 cells, we observed an increase in intracellular viral RNA starting at 4 h post infection, which continued to increase up to 24 h post infection (Fig. 1a-b). Increased extracellular virus RNA was observed starting at 6 h post infection which was paralleled by the release of infectious virus (Fig. 1a-c). The levels of viral RNA, production of infectious virus and virus spread were significantly higher in Calu-3 cells compared to A549-ACE2 cells (Fig. 1d-e).

**Fig. 1.**
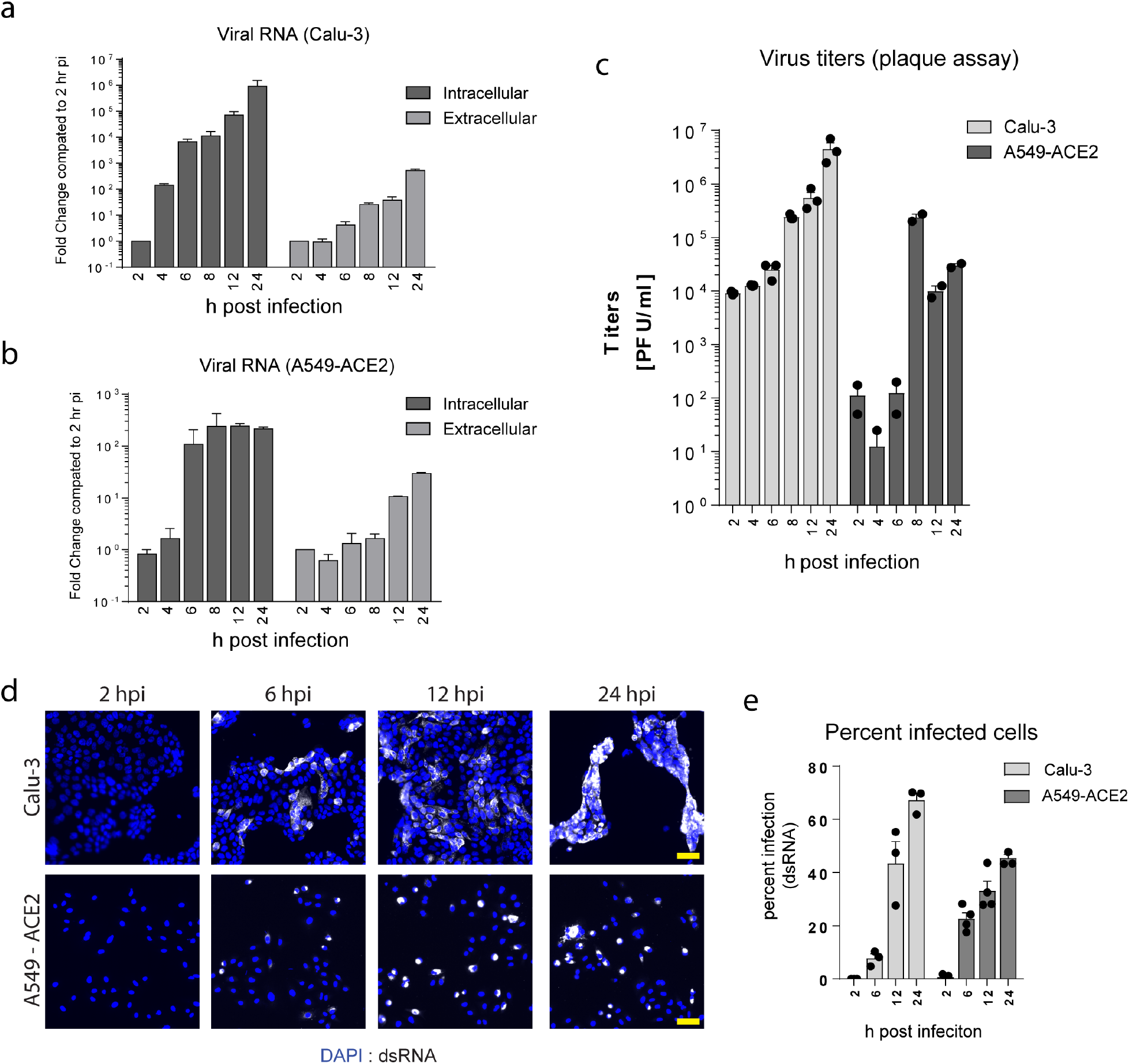
Virus growth kinetics in Calu-3 and A549-ACE2 cells. **a-e,** A549-ACE2 cells or Calu-3 cells were infected with SARS-CoV-2 (MOI=1). At the indicated times after infection samples were harvested for analysis of intra- and extra-cellular viral RNA (a-b), titer of infectious virus released into culture supernatant (c) and percent infected cells (d-e). **a-b,** Total cellular RNA was isolated from supernatants or cells at the given time points and virus RNA levels were determined using RT-qPCR with ORF1a specific primers detecting only the viral RNA genome, but not subgenomic mRNAs. Graphs show the average fold change and SEM for each time point compared to the 2 h time point. Intracellular viral RNA levels were corrected for total cell numbers using HPRT as a standard. **c**, Infectious virus titers were determined using a PFU assay; graph shows the mean and SEM PFU/mL for each time point. **d,** Infected cells were fixed in 4% PFA followed by staining with antibodies specific for dsRNA. Representative images of one out of 3 independent time series are shown. **e,** The percentage of infected cells at each time point was determined based on dsRNA staining. Graphs show the average percent infection and SEM over 3 independent experiments (N>1000 cells counted per experiment).

To determine the effects of SARS-CoV-2 infection on the host transcriptional architecture, total RNA was isolated at 4 h, 8 h, 12 h and 24 h post infection and analyzed by microarray (Fig. 2 and Extended Data Fig. 1). Analysis of significantly differentially expressed genes showed substantial transcriptomic changes in Calu-3 cells with a total of 3215 differentially expressed genes (FDR< 10%, Fig. 2a). Less genes were differentially expressed in A549-ACE2 cells (Fig. 2b). Principal component analyses (PCA) showed significant effects of SARS-CoV-2 infection on Calu-3 cells, especially at 24 h post infection (Extended Data Fig. 1a). We did not observe an overall decrease in total mRNA quality or large differences in probe intensity, thus showing no indication that SARS-CoV-2 infection causes a general transcriptional shutdown. Importantly, we observed a high degree of overlap between top significantly upregulated or downregulated genes from both cell lines (Extended Data Fig. 1b-c). However, in A549-ACE2 cells, there was less overall change in transcript levels following infection (Fig. 2b, and Extended Data Fig. 1a and 1d), which is likely due to the lower levels of infection (~40% vs 80% at 24 h of A549-ACE2 vs Calu-3, respectively)(Fig. 1e). Gene-set enrichment analysis of the transcriptional changes using curated “Hallmark” pathways showed a strong upregulation of inflammatory responses in both cell lines, with gene sets from NF-κB and IL6-STAT3 pathways showing a high degree of enrichment (Fig. 2c-e and Extended Data Fig. 1d)^38^. Interestingly, transcripts involved in the type I/III IFN pathways showed little change over the course of infection. These results suggest that in lung cells, the response to SARS-CoV-2 infection is dominated by pro-inflammatory, NF-κB-driven pathways, with little to no contribution of the antiviral IFN system.

**Fig. 2.**
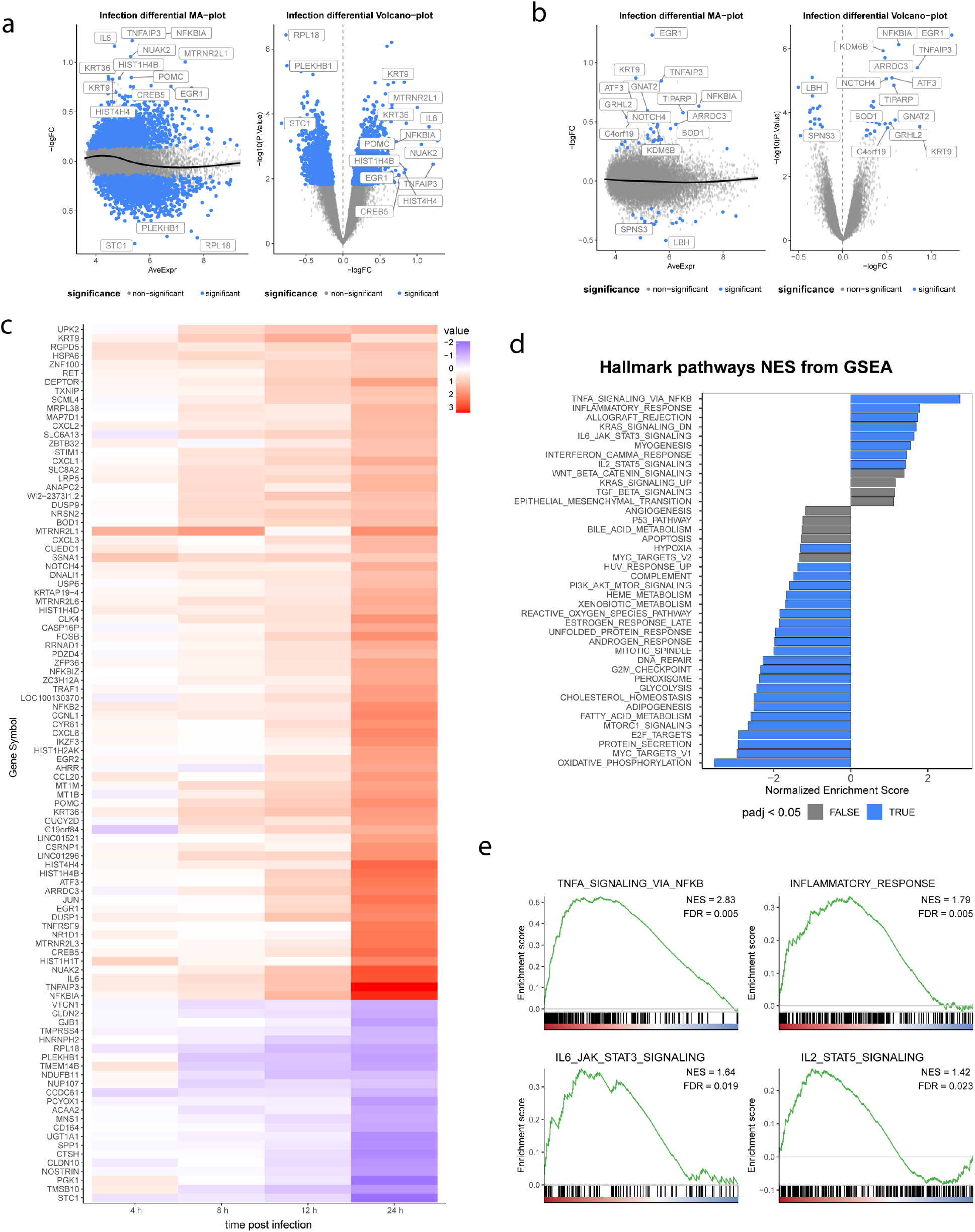
Transcriptional changes induced upon SARS-CoV-2 infection over time. **a-b,** A549-ACE2 cells or Calu-3 cells were infected with SARS-CoV-2. At the indicated time points after infection, total RNA was harvested and mRNA transcript levels were determined by Illumina microarray. Probe intensities were quantile normalized using probe-wise normalization. Normalized probe intensities were averaged for each gene and log-transformed. Differential expression was then called using a standard R/limma workflow for microarray analysis. MA-plots and Volcano-plots of transcriptional changes in Calu-3 cells (a) or A549-ACE2 cells (b) highlighting differentially expressed genes. Blue dots represent significant changes as determined by R/limma with a Benjamini Hochberg adjusted p-value smaller than 0.1. Left plot (MA-plot) shows log2 fold change on the y-axis vs mean normalized expression on the x-axis. Right plot (volcano-plot) shows the significant hits considering both infection and changes over time (x-axis = log2 fold change; y-axis = −log10 p-value; top 15 significant genes marked). **c.** Heat map of log scaled relative expression of enriched genes in Calu-3 cells. **d,** Gene set enrichment analysis employing the MSigDB collection of Hallmark pathways for Calu-3 cells. Top 40 enriched pathways for up- or down-regulated genes are shown, ranked by their normalized enrichment score. Color indicates significance (blue = adjusted p-value < 0.05) **e,** Barcode plots for pathways showing significant number of upregulated genes following SARS-CoV-2 infection in Calu-3 cells. NES = normalized area under the curve. FDR = Benjamini Hochberg adjusted p-value of enrichment (false discovery rate).

### Cell culture inflammatory responses parallels COVID-19 patient cytokine profiles

Transcriptional activation of NF-κB and inflammatory cytokine pathways in cultured cells infected with SARS-CoV-2 indicate that infected epithelial cells might contribute directly to the cytokine profiles observed in severe COVID-19 ^2^. To test this we compared the levels of secreted cytokines from infected Calu-3 cells to cytokine levels in serum, tracheal secretion, and bronchoalveolar lavage (BAL) fluid samples taken from infected patients with severe COVID-19 (Fig. 3a-b). Consistent with previous reports, infected patients had high levels of inflammatory cytokines including IL-6, TNF, Il-1β and IFNγ (Fig. 3a-d)^2^. Increases in IL-6 were also observed at early time points after infection of Calu-3 cells as well as high levels of the chemokine CXCL10/IP-10 at 24 h post infection (Fig. 3e). In contrast, no consistently detectable upregulation was found for IFNα, IFNβ, IFNγ and IFNλ2/3, whereas a rather moderate increase of IFNλ1 expression was found, but only at the late time point of Calu-3 cell infection. Together, these results corroborate published data showing that, in severe cases, SARS-CoV-2 infection preferentially induces a pro-inflammatory cytokine production with little activation of the antiviral responses. Additionally, these data indicate that infected epithelial cells secrete cytokines that can contribute to induction of tissue-level inflammation.

**Fig. 3.**
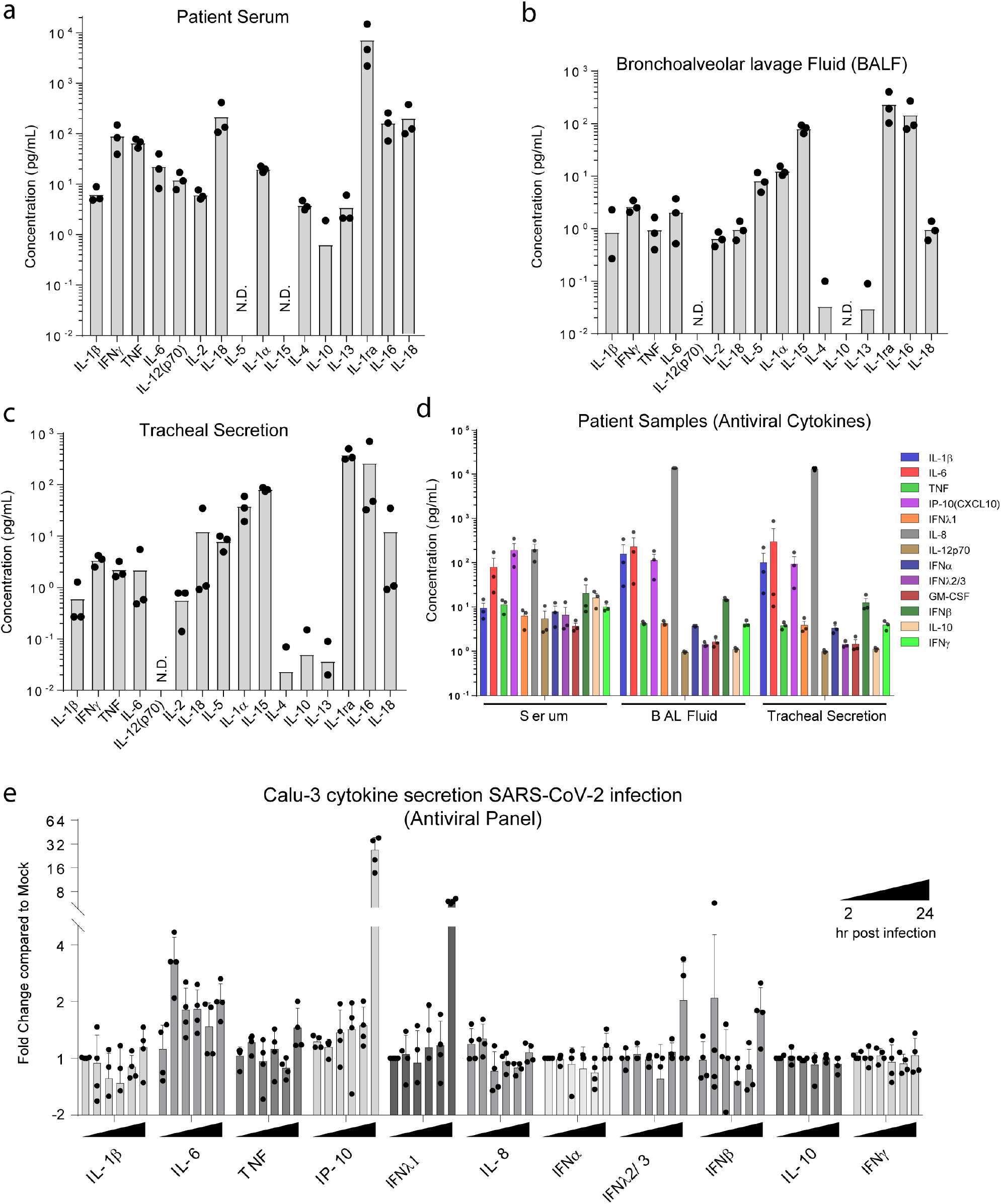
Cytokine profiles in various samples of SARS-CoV-2 infected patients and cells. **a-d,** Serum, bronchoalveolar lavage fluid, and tracheal secretion samples were taken from patients with severe COVID-19 symptoms. Samples were inactivated by treatment with beta-propiolactone that retains antigenicity. The cytokine profiles were determined using Extended Bio-Plex Pro Human Cytokine Panel (a-c) or LEGENDplex antiviral response panel (d) from LGENDplex. Graphs show the mean concentration for each cytokine (pg/mL) detected in the various patient samples. Each dot represents one patient sample. N.D., not detected. **e,** Calu-3 cells were infected with SARS-CoV-2. At 2 h, 4 h, 6 h, 8 h, 12 h, and 24 h post infection, cell culture supernatants were harvested from mock cells and infected cells, treated with betapropiolactone and cytokine profiles were determined by flow cytometry using the LGENDplex antiviral response panel. Values obtained for each infected sample were corrected for the corresponding value of the mock sample. Graph shows the mean and SEM of fold change for each cytokine compared to mock sample at each time point.

### SARS-CoV-2 infection is susceptible to type I/III IFN induced antiviral state

To confirm that the lack of IFN response in Calu-3 or A549-ACE2 cells infected with SARS-CoV-2 was not due to defects in the activation of innate immune pathways, we challenged these cells with dsRNA or other RNA virus infections. In Calu-3 cells transfected with dsRNA or infected with influenza virus or Sendai virus, we observed a robust upregulation of the IFN response (Fig. 4a). For A549 cells, similar transcriptional activation of ISGs was observed between wild type and ACE2-expressing cells in response to Zika virus (ZIKV) infection, a potent inducer of the IFN system (Fig. 4b-c). Notably, while magnitudes of the pro-inflammatory TNF mRNA were comparable between ZIKV and SARS-CoV-2 infection in A549-ACE2 cells, SARS-CoV-2 infection failed to induce IFN or classical ISGs, such as IFIT1 (Fig. 4c).

**Fig. 4.**
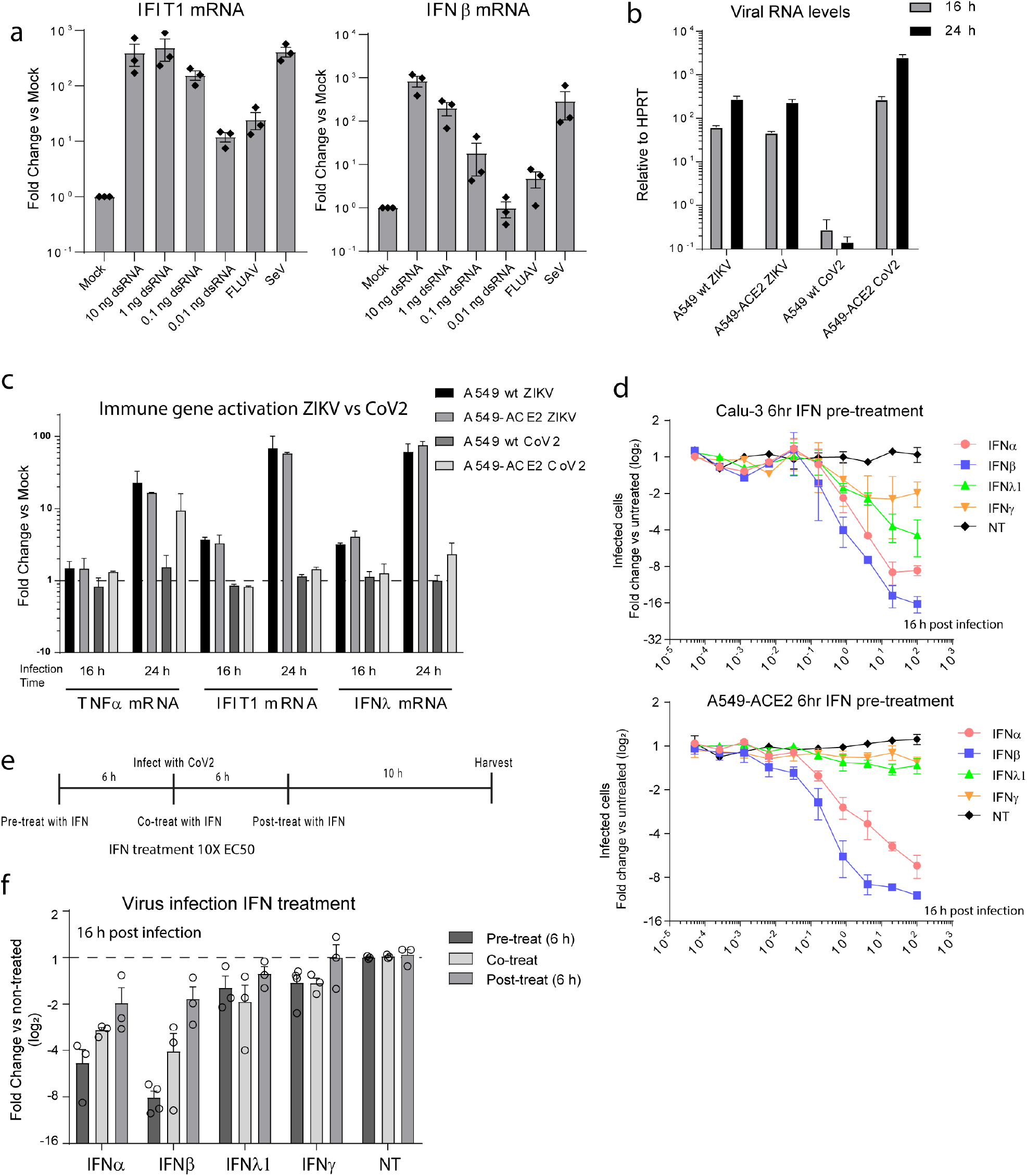
Innate cytokine response in lung epithelial cell lines and time-dependent inhibition of SARS-CoV-2 by IFN. **a,** Calu-3 cells were transfected with different concentrations of dsRNA or infected with influenza virus (FLUAV) or sendai virus (SeV) to activate the innate cytokine response. Total RNA was isolated from cells and the level of IFIT1 or IFNβ mRNA transcripts was determined by RT-qPCR. Graphs show the average and SEM for 3 independent experiments. **b-c**, Wild-type A549 or A549-ACE2 cells were infected with ZIKV or SARS-CoV-2 for the indicated time. Levels of viral RNA and immune gene transcripts were determined by RT-qPCR using specific primers. The graph in (b) shows the mean viral RNA levels compared to human HPRT levels for 3 independent experiments. The graph in (c) shows the mean fold change compared to uninfected cells for 3 independent experiments. **d,** Calu-3 or A549-ACE2 cells were treated with increasing concentrations of different IFNs starting with the highest concentration of 100X the EC50 value (determined for ISG activation by qPCR) for each IFN. 6 h post treatment cells were infected with SARS-CoV-2 (MOI=5) and 16 h thereafter cells were fixed in formaldehyde followed by staining with antibodies against dsRNA. The percent of infected cells was determined for each well. Graphs show the average fold change and SEM compared to untreated cells for 3 independent experiments. **e,** Experimental setup used for pre-, co-, and post-treatment of cells with IFN. **f,** Cells were treated with IFN for 6 h prior to infection (pre-treatment), or treated with IFN starting at the time of virus inoculation (co-treatment) or treated with IFN 6 h post infection (highest concentration of each IFN corresponds to 10X EC50 -determined for ISG activation by qPCR). 16 h post infection cells were fixed, and the percent infected cells was determined by immunofluorescence using dsRNA specific antibodies. Graph show the average fold change and SEM compared to untreated cells for 3 independent experiments.

Given the lack of an IFN response in SARS-CoV-2 infected cells, we next determined the effects of different IFN types on virus replication. Calu-3 or A549-ACE2 cells were pretreated with serial dilutions of type I, II or III IFNs, for 6 h followed by infection with SARS-CoV-2. In Calu-3 cells all IFNs reduced virus replication, with type I IFNs having the strongest effect, and IFNγ having the least effect (Fig. 4d – top panel; Extended Data Fig. 2). In the case of A549-ACE2 cells, IFNλ1 and IFNγ had little effect on virus replication, while IFNα or IFNβ pre-treatment robustly limited SARS-CoV-2 infection (Fig. 4d – bottom panel; Extended Data Fig. 2). These results are consistent with previous studies and further suggest that SARS-CoV-2 is susceptible to the antiviral state induced by IFNs prior to infection ^29,30^.

To test if IFNs could limit virus replication even after establishment of infection, A549-ACE2 cells were treated with high levels of various IFNs at the time point of infection or 6 h thereafter. Pre-treatment with type I IFNs, serving as control, blocked virus infection, whereas co- or post-treatment had significantly less effects (Fig. 4e-f). Of note, the 2-fold decrease in virus replication following post-treatment with type I IFNs likely represents a block in virus spread following the first round of infection, as only ~40-50% of cells were observed to be infected at the 6 h time point (Fig. 1e). Together, these observations suggest that SARS-CoV-2 likely supresses the production of IFNs and antiviral ISGs, and that it furthermore rapidly and potently blocks IFN signalling in infected cells.

### SARS-COV-2 infection specifically activates NF-κB but not IRF3

The high levels of inflammatory gene activation and the lack of IFNs and ISGs in response to SARS-CoV-2 infection lead us to investigate which transcription factors of the cell-intrinsic immunity are activated by the virus. In general, sensing of viral infection in epithelial cells by cytosolic innate immune receptors leads to the activation (i.e. phosphorylation) and nuclear translocation of the two hallmark transcription factors IRF3 and NF-κB, which in turn leads to transcriptional upregulation of antiviral and inflammatory genes. To evaluate the impact of SARS-CoV-2 infection on these pathways, we quantified the nuclear translocation of IRF3 and NF-κB in infected A549-ACE2 cells (Fig. 5a-c). Consistent with our transcriptomic data showing limited activation of antiviral genes, we observed no nuclear accumulation of IRF3 in SARS-CoV-2 infected cells, whereas a significant portion of infected cells showed nuclear translocation of NF-κB (Fig. 5a-b). In line with this, western blotting also showed increased levels of phosphorylated NF-κB p65/RELA starting at 12 h post infection accompanied by a decrease in IκB levels (Fig. 5d-e). These results suggest that in SARS-CoV-2 infection NF-κB is selectively activated or that IRF3 activation is supressed.

**Fig. 5.**
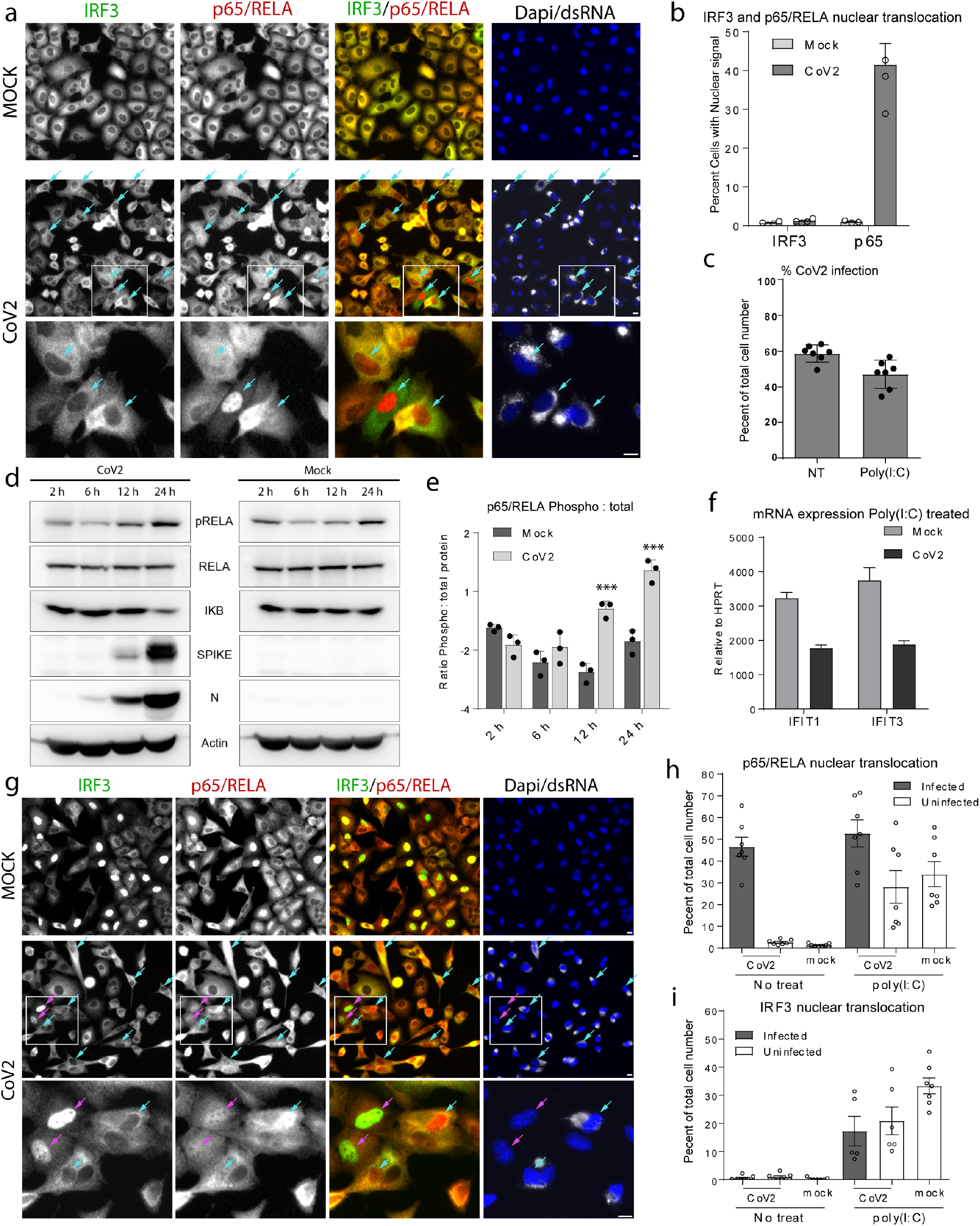
SARS-CoV-2 specifically activates the NF-κB pathway but not IFN/ISGs. **a-c,** A549-ACE2 cells were infected with SARS-CoV-2 for 16 h. **a,** Cells were fixed and stained with antibodies specific for IRF3 (green), p65/RELA (red) and dsRNA (grey). Turquoise arrows point to cells showing p65/RELA nuclear accumulation. Scale bars, 10 μm. **b,** Graph shows the mean nuclear accumulation of IRF3 and p65/RELA for images from cells treated as in panel (a). **c,** Infected cells we either untreated or transfected with Poly(I:C) 6 h post infection. Cells were fixed and stained with antibodies against dsRNA followed by analysis using widefield microscopy. Graph shows the average percent of infected cells for 7 fields of view collected from 3 independent experiments determined by dsRNA fluorescence signal (N>200 cells per field of view). **d-e,** Cells were infected with SARS-CoV-2 for the indicated times. Cells were lysed and levels of given proteins were determined by western blot using monospecific primary antibodies. **e,** Western blot signals for phosopho-p65/RELA (pRELA) were quantified and compared to the corresponding total p65/RELA proteins levels. Graph shows the mean and SEM for pRELA vs. total p65/RELA protein levels for 3 independent experiments. **f-i,** A549-ACE2 cells were infected with SARS-CoV-2 for 6 h followed by transfection with Poly(I:C) and incubated for 4 h. **f.** Total RNA was isolated and the mRNA levels of IFIT1 and IFIT3 were determined by RT-qPCR. Graphs show the mean and SEM from 3 independent experiments. **g,** Cells were fixed and stained with antibodies specific for IRF3 (green), p65/RELA (red) and dsRNA (grey). Magenta arrows point to cells with nuclear accumulation of both IRF3 and p65/RELA and turquoise arrows point to cells with only p65/RELA nuclear signal. Scale bars, 10 μm. **h-i,** Quantification of nuclear translocation of fluorescence signals from p65/RELA or IRF3 from 7 fields of view collected from 2 independent experiments conducted as in panel (g). Graphs show the mean number of cells with nuclear signal in uninfected or infected cells in either the Mock or SARS-CoV-2 treatment conditions. Infection was determined by the dsRNA signal. Quantification was done using an in house Fiji macro.

### SARS-CoV-2 interferes with poly(I:C)-mediated PRR activation

To determine if SARS-CoV-2 can actively block immune stimulation through cytosolic PRRs, 6 h after infection, cells were challenged by transfection with the dsRNA mimic poly(I:C). Activation of innate immune signaling was then assessed by monitoring the nuclear translocation of transcription factors and by measuring transcript levels of prototypic downstream ISGs. Compared to uninfected control cells, the upregulation of IFIT1 and IFIT3 in response to poly(I:C)-stimulation was significantly lower in SARS-CoV-2 infected cells (Fig. 5f). Consistently, we observed less nuclear translocation of IRF3 in response to poly(I:C) transfection in SARS-CoV-2 infected cells as compared to mock infected cells (Fig. 5g-i). Poly(I:C) transfection further lead to a slight reduction in the number of infected cells, likely due to limited virus spread (Fig. 5c – NT vs poly(I:C)). These data indicate a block of the PRR mediated IRF3 response in SARS-CoV-2 infected cells.

### Inflammatory response to SARS-CoV-2 infection is mediated by cGAS-STING

To determine the source of the SARS-CoV-2 induced inflammatory response or downstream immune activation, we evaluated the effects of innate immune receptor knockout or overexpression. We first looked at RNA receptors that have previously been described to recognize viral RNAs including RIG-I, MDA5 and TLR3 as well as IFN receptors. RIG-I/MDA5 double knockout, TLR3 overexpression, or IFN receptor (IFNAR, IFNGR, IFNLR) triple knockout A549-ACE2 cells were infected with SARS-CoV-2 and the upregulation of IFIT1 (IRF3 target) and TNF (NF-κB target) transcript levels were used as readouts for pathway activation. No significant changes in IFIT1 mRNA, TNF mRNA levels or viral RNA levels were observed in any of the cell lines compared to control cells (Extended Data Fig. 3a-b). Together, these data indicate that recognition of viral RNA via cellular RNA sensors is not involved in NF-κB activation in SARS-CoV-2 infected cells.

Although cGAS is a sensor of cytosolic DNA, induction of the cGAS-STING-signaling axis leading to activation of NF-κB and IRF3 has been reported for several RNA virus infections, most likely through cellular stress responses ^9^. To determine whether the cGAS-STING pathway is triggered in SARS-CoV-2 infection, we first evaluated changes in localization of cGAS or STING in infected cells. Indeed, both cGAS and STING were observed to re-localize to perinuclear clusters in infected cells, indicative of activation (Fig. 6a-b). Costaining for cGAS and dsDNA in infected cells also showed that dsDNA colocalized with cGAS in infected cells (Fig. 6c). Additionally, we observed that, unlike in poly(I:C)-mediated activation of RLRs, SARS-CoV-2 infection did not interfere with activation of the cGAS-STING pathway by dsDNA transfection (Extended Data Fig. 3c-d).

**Fig. 6.**
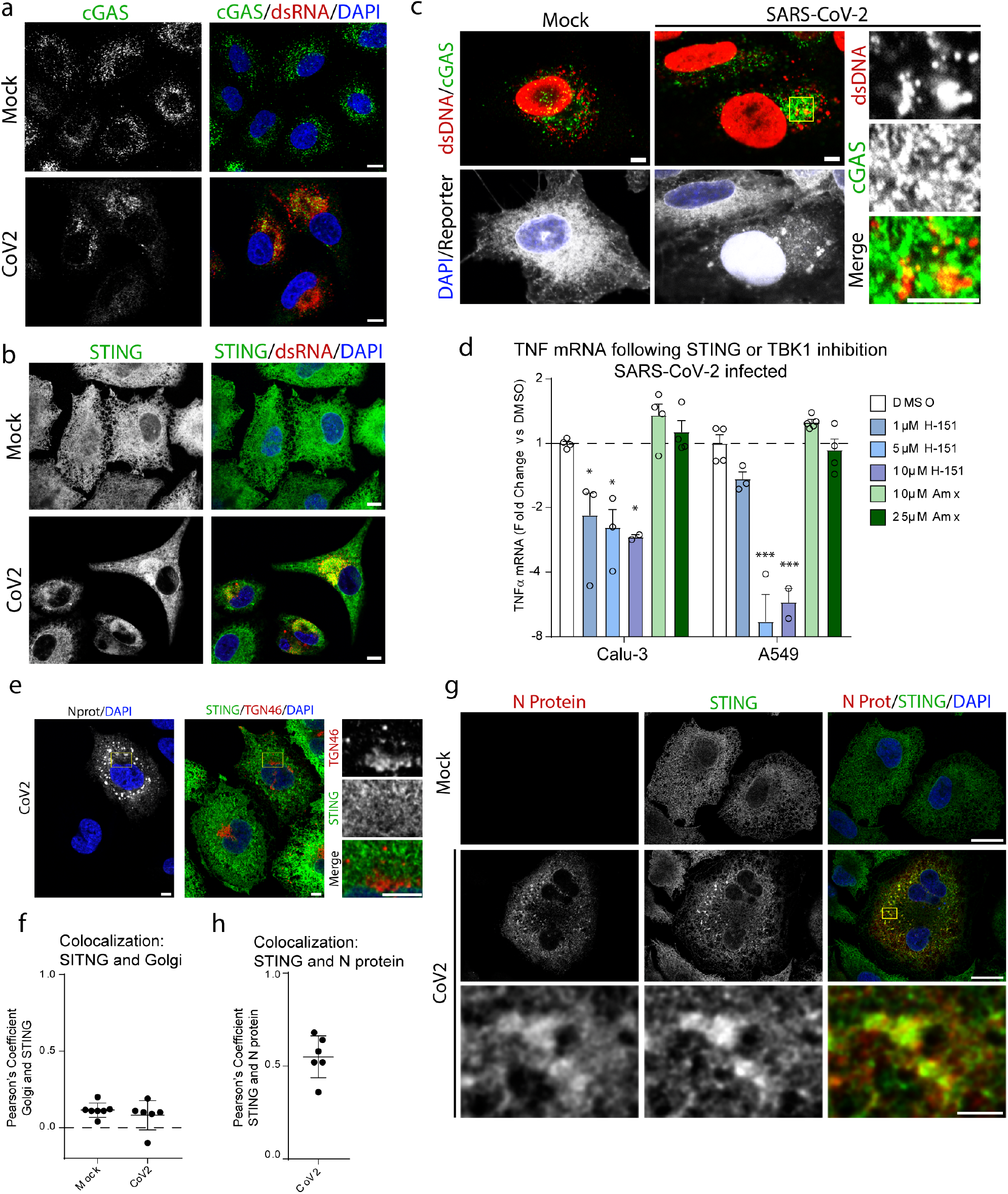
cGAS-STING activation mediates the NF-κB response in SARS-CoV-2 infected cells. **a-c,** A549-ACE2 cells were infected with SARS-CoV-2 for 16 h followed by fixation and staining with the indicated antibodies. Cells were analyzed by confocal microscopy. Scale bars 10 μm (a-b), 5 μm (c). **d,** Cells were infected with SARS-CoV-2. One hour after infection cells were treated with the indicated drugs at the given concentrations. Total RNA was isolated and the TNF mRNA transcript levels were determined by RT-qPCR. Graph shows the average fold change and SEM for TNF transcript levels compared to DMSO treated cells for 4 independent experiments. **e,** A549-ACE2 cells were infected with SARS-CoV-2 for 16 h followed by fixing and staining with the indicated antibodies. Cells were analyzed by confocal microscopy. Scale bars, 5 μm. **f,** Pearson’s correlation coefficient for fluorescence signals pertaining to STING and TGN46 were calculated for 7 fields of view over 2 independent experiments (N>20 cells). **g,** A549-ACE2 cells were infected SARS-CoV-2 for 16 h followed by fixing and staining with antibodies specific to STING (green) or N protein (red). Cells were analyzed by confocal microscopy. Scale bars, 10 μm upper panels, 1 μm for inset that is indicated with a rectangle in middle right panel. **h,** Pearson’s correlation coefficient for fluorescence signal pertaining to STING and N protein were calculated for 6 fields of view over 2 independent experiments (N>20 cells).

To confirm that cGAS-STING activation is involved in the observed induction of pro-inflammatory cytokines, we examined the effects of pharmacologically blocking STING in SARS-CoV-2 infected cells. One hour post infection, cells were treated with the STING specific inhibitor H-151, the TBK1 inhibitor amlexanox (Amx), or DMSO. At 24 h post infection, we observed a significant decrease in the levels of TNF mRNA in infected cells treated with H-151 compared to DMSO treated cells (Fig. 6d), both, in A549-ACE2 and Calu-3. This decrease was not observed for Amx treated cells. Infection levels and cell viability were not significantly affected at the effective concentration (Extended Data Fig. 3e-g). Together these results indicate that SARS-CoV2-infection triggers the cGAS-STING pathway, leading to NF-κB-mediated induction of pro-inflammatory cytokines, and that this response can be controlled with STING inhibitors.

Although STING activation is usually associated with both, NF-κB as well as IRF3 activation, several reports have suggested that interfering with proper translocation of STING from the ER to Golgi compartments can selectively stimulate the NF-κB pathway ^39,40^. To test whether this is the case in SARS-CoV-2 infected cells, we determined the localization of STING relative to Golgi markers by microscopy. Consistent with previous reports, in cells transfected with dsDNA, we observed STING translocation to the Golgi compartment (Extended Data Fig. 3h). No significant colocalization of Golgi markers and STING were observed in either mock or SARS-CoV-2 infected cells, suggesting that STING translocation may be impaired (Fig. 6e-f). Moreover, we found that clusters of STING in SARS-CoV-2 infected cells colocalized with viral nucleocapsid (N) protein (Fig. 6g-h; Extended Data Fig. 3i). Together, these results suggest that STING is activated in SARS-CoV-2 infection, but inhibited from translocating to the Golgi, leading to a specific NF-κB inflammatory response in infected cells.

## Discussion

In this study, we combine transcriptional profiling and cytokine secretion analyses to characterize the pro-inflammatory response induced by SARS-CoV-2 infection, and evaluate the virus-induced signalling pathways mediating this response. We report that both virus-induced transcriptional changes and cytokine profiles from infected epithelial cell cultures parallel cytokine profiles from patient BAL fluid, tracheal secretions and serum samples, indicating a role for infected epithelial cells to contribute to the hyper-inflammatory response described for patients suffering severe COVID-19. This pro-inflammatory response in infected epithelial cells is initiated by an activation of NF-κB with a concurrent robust block of the IRF3 and IFN pathways. We further demonstrate that this activation of NF-κB is not mediated by the expected viral RNA recognizing receptors of the RLR or TLR family, but instead SARS-CoV2 infection leads to the activation of the cGAS-STING signaling axis. Putatively by preventing relocalization of activated STING to the Golgi, SARS-CoV-2 appears to specifically allow for NF-κB induction while omitting the activation of the antiviral IRF3 / IFN system. Intriguingly, upregulation of NF-κB-regulated pro-inflammatory cytokines, such as TNF, can be efficiently blocked by the administration of pharmacological STING inhibitors.

Building on recent reports, our results demonstrate that SARS-CoV-2 infection causes an imbalanced immune response in infected lung epithelial cells, resulting in an NF-κB polarized response rather than a classic antiviral immune response (NF-κB, IRF3/7 and IFN signalling), which is likely amplified by immune cells to produce the cytokine storm symptoms associated with COVID-19 ^3,29,41,42^. Pro-inflammatory cytokine responses are an important aspect of the innate immune response that are required to recruit professional immune cells to the site of infection and aid in the initiation of the adaptive immune response. This response, together with the activation of antiviral pathways, including type I/III IFNs, creates a potent antiviral environment. Our study, in combination with several parallel studies indicates that SARS-CoV-2 infection induces a selective inflammatory response that can cause pathogenic inflammation without effectively controlling the virus. Similar inflammatory activation has been observed for SARS-CoV-1 infections, where the viral ORF3a activates NF-κB to trigger the NLRP3 inflammasome ^43^. Additionally, NF-κB inhibition can increase mouse survival in SARS-CoV-1 infected mice ^44^. We note that several studies have indicated that SARS-CoV-2 infection leads to IFN production in lung or intestinal tissue ^30,33^. Our microarray results show only a slight increase in the IFNλ1 response, and only at late time points after infection, which indicates that the efficient inhibition of the IRF3 signalling pathway might be bypassed in some cells with heavy virus load. Increased immune activation has also been reported when cells are infected with excessive amounts of virus ^29^. This could be due to high levels of cellular stress or high levels of virus dsRNA in cells at these later time points. In either case, this immune activation indicates that SARS-CoV-2 infection does not completely block the antiviral immune activation, at least at late time points after infection.

We demonstrate that type I/III IFNs block SARS-CoV-2 infection in lung cells, consistent with several recent studies, but this effect is rapidly diminished once virus replication has been established. These results corroborate the notion that SARS-CoV-2 infection leads to a robust block in IFN receptor mediated signaling, which is similar to observations with SARS-CoV-1 and MERS-CoV. Additionally, our transcriptional analysis and cytokine profiles show that SARS-CoV-2 infection also fails to produce a significant IFN response in infected lung epithelial cells, which could trigger a paracrine response, limiting virus spread. In addition to blocking IFN signalling, SARS-CoV-2 infection limits the activation of dsRNA sensing PRR pathways even when exogenously stimulated by poly(I:C) transfection. Together these results demonstrate that SARS-CoV-2 infection, similar to SARS-CoV-1 and MERS-CoV, employs mechanisms to robustly block immune sensing of viral RNA by RLRs, as well as the downstream immune signalling pathways ^17,18^.

Further evaluation of the SARS-CoV-2 induced pro-inflammatory response showed a specific induction of NF-κB, but not of IRF3 or the subsequent IFN signalling. NF-κB can be activated through numerous immune or stress stimuli including the ER stress responses or increase in cytosolic reactive oxygen species, as well as through detection of cytosolic DNA released from the nucleus or mitochondria (reviewed in ^8,45,46^). Our results indicate that cGAS-STING activation is a major contributor to NF-κB activation in SARS-CoV-2 infected cells. Since cGAS is a dsDNA sensor that would not be expected to directly recognize SARS-CoV-2 RNA, it is likely that cellular stress or cytokine responses induced by the infection leads to nuclear or mitochondrial DNA release which is sensed by cGAS^47-51^. Similar activation of cGAS-STING has been observed for other positive strand RNA viruses including flaviviruses and both SARS-CoV-1 and NL63 coronaviruses (reviewed in ^9^). For the coronaviruses, STING activation is perturbed through the action of the viral PLP leading to an inhibition of STING oligomerization and downstream activation of TBK1 and IRF3 ^28,52,53^. Intriguingly, mechanisms for cGAS-STING modulation seem to be different in SARS-CoV-2 infected cells, highlighting a major immunological difference between these related viruses. Of note, the activation of NF-κB through cGAS-STING does not exclude other sources of NF-κB activation. Indeed, we observed increases in FOS/JUN and ATF3 mRNA levels in infected cells suggesting activation of multiple cell stress pathways ^54^. Moreover, pharmacological inhibition of STING did not completely block TNF upregulation, further indicating a role for other sources of NF-κB-activation. We speculate that therapeutic inhibition of multiple NF-κB activation pathways could serve to further reduce pro-inflammatory responses in SARS-CoV-2 infected cells.

The selective activation of NF-κB, rather than a general block in all immune activation pathways, indicates a pro-viral role for NF-κB signalling. In addition to functions in inflammation, NF-κB is also important for cell survival and proliferation ^55^. These NF-κB cell survival signals could be beneficial for the virus by promoting vitality in cells in order to facilitate efficient and sustained virus replication and spread. Mechanisms for NF-κB pathway interference have been reported for numerous DNA and RNA viruses ^56–58^. Selective modulation of the cGAS-STING pathways may allow SARS-CoV-2 to promote an NF-κB mediated cell survival signal while limiting ISG induction.

Classical cGAS-STING induction activates not only NF-κB, but also TBK1 and IRF3 pathways. We envisage several mechanisms that could contribute to the selective NF-κB activation. First, the virus could actively block TBK1 activation in infected cells. Indeed, protein interaction studies indicate that viral NSP13 and NSP15 proteins interact with TBK1 or its adaptor proteins ^34^. Additionally, a block in TBK1 activation has been reported for both SARS-CoV-1 and MERS-CoV infections and early reports demonstrate a lack of TBK1 phosphorylation in SARS-CoV-2 infected cells ^29,59^. Moreover, our results support a model where SARS-CoV-2 infection prevents activated STING from translocating from the ER to the Golgi. Activation of STING at the ER has been shown to be sufficient for NF-κB activation but not for TBK1 activation and the subsequent IRF3 phosphorylation ^40,60^. It may be that fragmentation of the Golgi by SARS-CoV-2 infection leads to an impairment of STING translocation to the ERGIC. Consistently, our transcriptomic analysis shows impairment in protein secretion pathways, specifically including downregulation of several COP coatomer proteins involved in ER to Golgi transport. Alternatively, SARS-CoV-2 proteins could actively block cGAS-STING translocation. Colocalization between STING and N protein in infected cells suggests a direct role for N protein in limiting STING translocation. A similar mechanism has been suggested for murine cytomegalovirus, where viral m152 protein associates with STING and limits exit from the ER, thereby promoting an NF-κB specific response. Interestingly, pathway analysis of our microarray data indicate that cytokine transcriptional responses from SARS-CoV-2 infected cells resemble signatures from human cytomegalovirus infected cells. Further experimentation is required to define the precise mechanisms of STING activation in SARS-CoV-2 infected cells and to determine whether viral proteins are directly associating with components of the cGAS-STING pathway.

The majority of documented SARS-CoV-2 infections lead to mild or no symptoms, indicating that even the observed low level of antiviral pathway activation induced by infected cells can be sufficient to limit and resolve the infection. On the other hand, in patients with underlying conditions or attenuated immune responses, these antiviral responses do not limit virus replication and a sustained virus load eventually leads to a long term inflammatory response. In these latter cases, one important avenue of treatment is to modulate the immune response in order to alleviate hyper-inflammation. In addition to other immune modulators that are currently being used or clinically evaluated (eg. IL-6 inhibitors or corticosteroids)^61–65^, our results indicate that disease severity might be suppressed at the epithelial cell level through the use of cGAS-STING inhibitors or through blocking NF-κB mediated inflammatory responses. In this respect, NF-κB inhibitors analogous to CAPE or parthenolide, prolonging survival of SARS-CoV-1 infected mice ^44^, might help to reduce the disease burden imposed by COVID-19.

## Methods

### Cell lines and culture conditions and viruses

A549 and Calu-3 cells were cultured in Dulbecco’s Modified Eagle’s Medium (DMEM) supplemented with Glutamax (Gibco),10% fetal bovine serum, 100 U penicillin/ml, 100 μg streptomycin/ml, 2 mM L-glutamine and nonessential amino acids. A549 cells stably expressing ACE2 and the SARS-CoV-2 reporter construct were created by lentivirus transduction. To produce lentivirus particles, HEK-293T cells were transfected with pCMV-Gag-Pol, pMD2-VSV-G (kind gifts from Didier Trono, EPFL, Lausanne, Switzerland) and a pWPI vector encoding the gene of interest. Transfections were done using polyethylenimine and lentivirus particles were harvested and filtered through a 0.45 μm pore-size filter. A549 cells were inoculated with the viral supernatant overnight and next day antibiotic selections were applied. Neomycin (500 μg/ml) and Puromycin (2 μg/ml) antibiotics were used for ACE2 and SARS-CoV-2 reporter expressions, respectively. Viruses used are SARS-CoV-2-BavPat1/2020 strain (kindly provided by Christian Drosten through the European Virus Archive), ZIKV H/PF/2013 (GenBank accession number KJ776791.2), IAV A/WSN/1933 (H1N1; kindly provided by Martin Schwemmle, University of Freiburg) and SeV (kindly provided by Rainer Zawatzky, German Cancer Research Center).

### SARS-CoV-2 virus stock production

SARS-CoV-2 stocks were produced using VeroE6 cell line. Passage 2 BavPat1/2020 (MOI: 0.01) strain was used to generate the seed virus (passage 3). After 48 h the supernatant was harvested, cell debris was removed by centrifugation at 1,000xg for 5 min and supernatant filtered with a 0.45 mm pore-size filter. Passage 4 virus stocks were produced by using 500 μl of the seed virus (passage 3) to infect 9E+06 VeroE6 cells. The resulting supernatant was harvested, filtered 48 h later as described above and stored in aliquots at −80°C. Stock virus titers were determined by plaque assay.

### RNA isolation and RT-qPCR

Total RNA was isolated from cells or supernatants using the NucleoSpin RNA extraction kit (Macherey-Nagel) according to the manufacturer’s specification. cDNA was synthesized from the total RNA using the high capacity cDNA reverse transcription (RT) kit (ThermoScientific) according to the manufacturer’s specifications. Each cDNA sample was diluted 1:15 in nuclease free H2O prior to qPCR analysis using specific primers and the iTaq Universal SYBR green mastermix (Bio-Rad). Primers for qPCR were designed using Primer3 software and include: SARS-CoV-2-ORF1 fwrd-5’-GAGAGCCTTGTCCCTGGTTT-3’, rev-5’-AGTCTCCAAAGCCACGTACG-3’; IFIT1 fwrd-5’-GAAGCAGGCAATCACAGAAA-3’, rev-5’-TGAAACCGACCATAGTGGAA-3’; IFIT3 fwrd-5’-GAACATGCTGACCAAGCAG-3’, rev-5’-CAGTTGTGTCCACCCTTCC-3’; TNF fwrd-5’-TAGCCCATGTTGTAGCAAACCC-3’, rev-5’-GGACCTGGGAGTAGATGAGGT-3’ GAPDH fwrd-5’-GAAGGTGAAGGTCGGAGTC-3’, rev-5’-GAAGATGGTGATGGGATTTC-3’; HPRT fwrd-5’-CCTGGCGTCGTGATTAGTG-3’, rev-5’-ACACCCTTTCCAAATCCTCAG-3’. To obtain the relative abundance of specific RNAs from each sample, cycle threshold (ct) values were corrected for the PCR efficiency of the specific primer set, and normalized to hypoxanthine phosphoribosyltransferase 1 (HPRT) transcript levels.

For microarray chip analysis total RNA was extracted from cells and hybridized on an Affymetrix Clariom S human array performed by the Microarray Unit of the Genomics and Proteomics Core Facility at the German Cancer Research Center (DKFZ). Labeling was done using the Thermo Fisher Scientific (Affymetrix) Gene Chip WT PLUS Reagent to generate labeled ss-cDNA from input amounts of 50 ng total RNA. Hybridization was done according to manufacturer’s protocol for Thermo Fisher Scientific (Affymetrix) Gene Chip WT PLUS Reagent Kit. 5.5 μg of fragmented and labeled ss-cDNA were hybridized for 17 hr at 45°C on Thermo Fisher Scientific (Affymetrix) human Clariom S Arrays. Chip scanning Gene Expression Microarrays were scanned using the Affymetrix GeneChip® Scanner 3000 according to GeneChip® Expression Wash, Stain and Scan Manual for Cartridge Arrays

### Data analysis for microarray

Raw, analyzed and meta data as well as the code used during analysis are available upon request.

First, data was collected for all samples after Robust Multi-Array Average (RMA) quantile normalization with R using the function ‘normalize.quantiles’ from Bioconductor package “preprocessCore” for probe set equalization. Second, data was then log transformed and PCA including all samples was performed using R/prcomp (R version 4.0.0). The rotation for each sample is shown. After PCA quality control and check for equal distribution of log-transformed probe intensities, data was gathered and time points were pooled as ‘early’ (4h and 8 h time point) or ‘late’ (12 h and 24 h). R/limma’s lmFit, eBayes and topTable functions were then used with a model “matrix of expression ~ treatment + time” (limma version 3.40.6,^66^), to estimate base mean expression and differential expression for the contrast infected vs mock treatment. This analysis was performed individually for each cell line as differences between the lines would have obscured a model by driving the variance, as apparent in the PCA analysis. R/limma’s topTable function employs Benjamini Hochberg correction for multiple testing on all p-values. Genes were called significant if their adjusted p-value was smaller than 0.1 (False Discovery Rate, FDR < 10%).

Gene set enrichment analysis was performed according to Subramanian et al. ^67^. We use the practical R implementation “fgsea” ^68^ and the hallmark pathway gene set published by Liberzon et al. ^69^. The barcode plot implementation was inspired by Zhan et al.^70^.

### Antibodies

Primary antibodies and specific dilutions used for western blot or immunofluorescence included: Mouse anti-dsRNA J2 (Scicons: 10010500, IF-1:1000); Mouse anti-SARS-CoV-2 N protein (Sino Biological: 40143-MM05, IF - 1:1000; WB - 1:1000); Rabbit anti-SARS-CoV-2 Spike protein (Abcam: ab252690, WB-1:1000); Rabbit anti-IRF3 (Cell Signaling Technology: 11904S, IF - 1:400); Mouse anti-P65/RELA (Santa Cruz: sc-8008, IF – 1:100); Rabbit anti-cGAS (Atlas Antibodies: HPA031700, IF – 1:100); Rabbit anti-STING (Atlas Antibodies: HPA038534, IF – 1:100); Mouse anti-dsDNA (Abcam: ab27156, IF – 1:2000); Rabbit anti-p65/RELA (Cell Signaling: L8F6, WB – 1:1000); Rabbit anti-phospho-p65/RELA (Cell Signaling: 3033, WB - 1:1000); Rabbit anti-IkB (Cell Signaling: 9242s, WB - 1:1000); Sheep and-TGN46 (Biorad:AHP500G, IF-1:200); Mouse anti-Actin (Sigma Aldrich: A5441, WB-1:5000).

Secondary antibodies used for western blot included Goat anti–rabbit IgG-HRP (Sigma Aldrich A6154, 1:2000), Goat anti–mouse IgG-HRP (Sigma Aldrich A4416, 1:5000). Secondary antibodies for immunofluorescence included: Alexa Fluor 488 donkey anti-rabbit IgG (Thermofisher A-21206), Alexa Fluor 488 donkey anti-mouse IgG (Thermofisher A-21202), Alexa Fluor 488 donkey anti-mouse IgG2a (Thermofisher A-21131), Alexa Fluor 568 donkey anti-rabbit IgG (Thermofisher A-10042), Alexa Fluor 568 donkey anti-mouse IgG (Thermofisher A-10037), Alexa Fluor 568 donkey anti-mouse IgG1 (Thermofisher A-21124), Alexa Fluor 647 donkey anti-rabbit IgG (Thermofisher A-31573), Alexa Fluor 647 donkey anti-mouse IgG (Thermofisher A-31571). ALL Alexa fluor secondary antibodies were used at 1:1000.

### Immunofluorescence analysis

After infection with SARS-CoV-2 cells were fixed with 6% formaldehyde solution, washed twice with phosphate buffered saline (PBS) and permeabilized with 0.2% Triton X-100 in PBS. Next, the Triton X-100 solution was replaced with 2.5% (w/v) milk solution (in PBS) and cells were blocked for 1 h at room temperature. Primary antibodies were diluted in 2.5% milk solution and samples were incubated with primary antibodies for 1 h. After washing three times with PBS, samples were incubated with Fluorophore-conjugated secondary antibodies, diluted in milk solution, for 30 min. After washing three times with PBS samples were mounted in Fluoromount G solution containing DAPI (Southern biotech) for DNA staining. Microscopic analyses were conducted with a Nikon Eclipse Ti microscope (Nikon, Tokio, Japan) or a Leica SP8 confocal microscope (Leica) for the subcellular localization analyses.

For quantification of the nuclear translocation of NF-κB p65/RELA or IRF3, nuclei were segmented using DAPI signal first. Secondly, the segmented nucleus was dilated and finally dilated nucleus was subtracted by original nucleus mask to detect perinuclear fluorescent signal. To determine SARS-CoV-2 infected cells, dsRNA intensity was measured within the perinuclear area. The status of NF-κB or IRF3 nuclear signals was determined based on the ration between perinuclear intensity divided by nuclear intensity.

### Interferon treatment

For pretreatment experiments, cells were treated for 6 h with serial dilution of IFNα2 (PBL Assay Science, 11100-1), IFNβ (R&D Systems, 8499-IF-010/CF), IFNλ1 (Peprotech, 300-02L-100) or IFNγ (R&D Systems, 285-IF/CF) in normal cell growth media. The 50 percent effective concentration (EC50) values for ISG activation were determined using RT-qPCR or from the data sheet (for IFNγ) and IFN dilutions were started at 100 times the EC50 value. For pre-, co-, and post-treatment with IFNs, cells were treated with each IFN at 10 times the EC50 concentration of each IFN. For all IFN treatments, cells were fixed with 4% paraformaldehyde at 16 h post infection, followed by immunofluorescence staining as describe above. Cells were imaged using a Nikon Eclipse Ti2-E inverted fluorescent microscope. In order to determine the percentage of infected cells, we first segmented single cells, and secondly, calculated the mean signal intensity value of dsRNA from SARS-CoV-2 located in the perinuclear region of segmented single cells. Fluorescent signal intensity values of dsRNA were quantified using the EBImage platform in R-Studio^71^. Finally, percent infected cells was determined by establishing a threshold for the background fluorescent signal intensity of non-infected cells and quantifying the number of cells that exceeded this threshold in a given field of view.

### Poly(I:C) and herring DNA transfection

For Calu-3 cell stimulation (Fig. 4a,b) cells were transfected with the indicated amount of poly(I:C) using lipofectamine 2000 reagent as per the manufacturer’s protocol. 16 h after transfection, total RNA was isolated and RT-qPCR was used to determine transcript levels as describe above.

For transfection in SARS-CoV-2 infected cells (Fig. 5 and extended data Fig. 3), cells seeded in 24-well plates were infected with SARS-CoV-2 at MOI=5 for 6 h. Cells were then transfected with poly(I:C) or herring DNA (500ng/well) using lipofectamine 2000 reagent as per the manufacturer’s protocol. 6 h after transfection, cells were either fixed with 4% paraformaldehyde and processed for immunofluorescence, or total RNA was isolated for RT-qPCR analysis as described above.

### Western blot analysis and imaging

Infected and mock cells were washed with PBS and lysed with 100 μl of sample buffer (120 mM Tris-HCl [pH 6.8], 60 mM SDS, 100 mM DTT, 1.75% glycerol, 0.1% bromophenol blue) supplied with 1 μl of benzonase (Milipore: 70746-3) to remove contaminating nucleic acids. Denaturation of the samples was achieved by incubation at 95°C for 3 min. After SDS-PAGE, proteins were blotted onto PVDF (polyvinylidenfluorid) membranes and blocking was done with 3% (w/v) BSA in Tris-buffered saline (TBS) for 1 h at room temperature. Membranes were incubated with primary antibodies (Extended Data Table 1), diluted in 3% BSA in TBS, for 1 h and washed three times for 10 min each with TBS-T (TBS supplied with 0.1% Tween 20). Horse radish peroxidase (HRP)-conjugated secondary antibodies were diluted in 5% (w/v) milk in TBS-T and membranes were incubated for 1 h at room temperature. After washing three times with TBS-T for 10 min, membranes were developed with the Western Lightning Plus-ECL reagent (Perkin Elmer: NEL105001EA). A ChemoCam Imager 3.2 (Intas Science Imaging Instruments GmbH, Göttingen, Germany) was used to visualize the signals that were quantified using the ImageJ (FiJi) software package.

### Plaque assay and CPE assay

2.5E+06 VeroE6 cells were seeded into each well of a 24-well plate. On the next day, cells were infected with serial dilutions of SARS-CoV-2 for 1 h. Afterwards inoculum was removed and replaced with serum-free DMEM containing 0.8% carboxymethylcellulose. At 72 h post infection, cells were fixed with 5% formaldehyde for 1 h followed by staining with 1% crystal violet solution. Plaque forming units per ml (PFU/ml) were calculated by manual counting of the viral plaques.

For cytopathic effect (CPE) assays, Calu-3 or A549-ACE2 cells were plated into 96 well plates. Cells were infected with SARS-CoV-2 for 48 h followed by fixation in 5% formaldehyde for 1 h. Cells were then stained with 1% crystal violet solution and scanned.

### Cytokine profiling of patient samples

Patient sera were collected and stored at −80°C until cytokine measurement. Bronchoalveolar lavage fluid BALF specimens and tracheal secretions (TS) were collected during medically-indicated, routine bronchoscopies in mechanically ventilated COVID-19 patients using a flexible fiberoptic bronchoscope. BAL-samples were obtained by instillation of a pre-warmed 0.9% sterile saline solution (20 mL twice). The sampling area was determined based on the localization of lesion on chest imaging (X-ray or computed tomography scan). All material was obtained after approval by the by the Ethics Committee of the Medical Faculty Heidelberg (number S-148/2020) medical ethics committee of the University of Heidelberg, written consent was obtained from all patients prior to analysis.

Blood, BALF and TS samples were evaluated for cytokine for cytokine levels and compared to the cytokine profiles secreted from infected culture cells. Serum was separated from clotted blood fraction by centrifugation at 1500xg for 10 min at 4°C. BALF and TS were filtered through a 100 μm cell strainer (Falcon Corning, USA) then centrifuged at 500xg 10 min at 4°C to remove cells and supernatants were treated with 0.05% beta-propiolactone (BPL) overnight, followed by a 2 h incubation at 37°C. Proteins contained in BALF and TS were concentrated 5x using an Amicon Ultra column 100kDa (Merck Millipore) and 20 min centrifugation at 4000xg and samples were stored at −80°C. Analyses were performed with the Extended Bio-Plex Pro Human Cytokine 48-Plex Screening Panel (Biorad, Munich, Germany) as described previously ^72^ and using a 2-laser reader allowing simultaneous quantification of cytokines and chemokines. Standard curves and concentrations were calculated with the Bio-Plex Manager software using the 5-parameter logistic plot regression formula. The detection sensitivity of all analyses ranged from 2 pg/mL to 30 ng/mL. Alternatively, samples were analysed by flow cytometry (BD LSRFortessa from BD Biosciences) using LEGENDplex antiviral response panel (BioLegend) according to the manufacturer’s instruction. All material was obtained after approval by the medical ethics committee of the University of Heidelberg; written consent was obtained from all patients prior to analysis.

## Acknowledgments

We thank all members of the Molecular Virology department at Heidelberg University for helpful discussions and support during different stages of the COVID-19 related lockdown. We also thank Dr. Monica Boxberger for taking patient samples, Sandra Wüst for excellent technical assistance, the Infectious Diseases Imaging Platform (IDIP) for facility use and help with microscopy, the imaging facility of the German Center for Infection Research (DZIF), and the Microarray Unit of the Genomics and Proteomics Core Facility at the German Cancer Research Center (DKFZ), for providing excellent expression profiling services.

Work by R.B. was supported by DZIF, project numbers 8029801806 and 8029705705, and the Deutsche Forschungsgemeinschaft (DFG, German Research Foundation) – project number 240245660 – SFB 1129 and project number 272983813 – TRR 179. Work in the laboratory of M. Bo. was, in part, supported by the ERC-Synergy Grant DECODE.

## Author contributions

This project was designed by C.J.N., B.C., M.C., M.Bi.. and R.B.. Molecular cloning, virus infection and cell culture treatment experiments were carried out by C.J.N, B.C. and M.C.. Microarray data analysis was done by F.H.. Culture cell line testing for immunocompetence was done by S.J., S.S.B., and D.Y.Z.. Microscopy image acquisition was done by C.J.N. and J.F. and image analysis was done by C.J.N., J.F., J.-Y.L., M.G., and B.E.-D.. Patient samples were collected by U.M.. Cytokine secretion evaluation and analysis was done by A.P. and N.H.. The manuscript was prepared by C.J.N, B.C., J.F., M.Bi., and R.B.. The work was supervised by M.Bo., M.Bi., and R.B..

## Extended Data Figures

**Extended Data Fig. 1.**
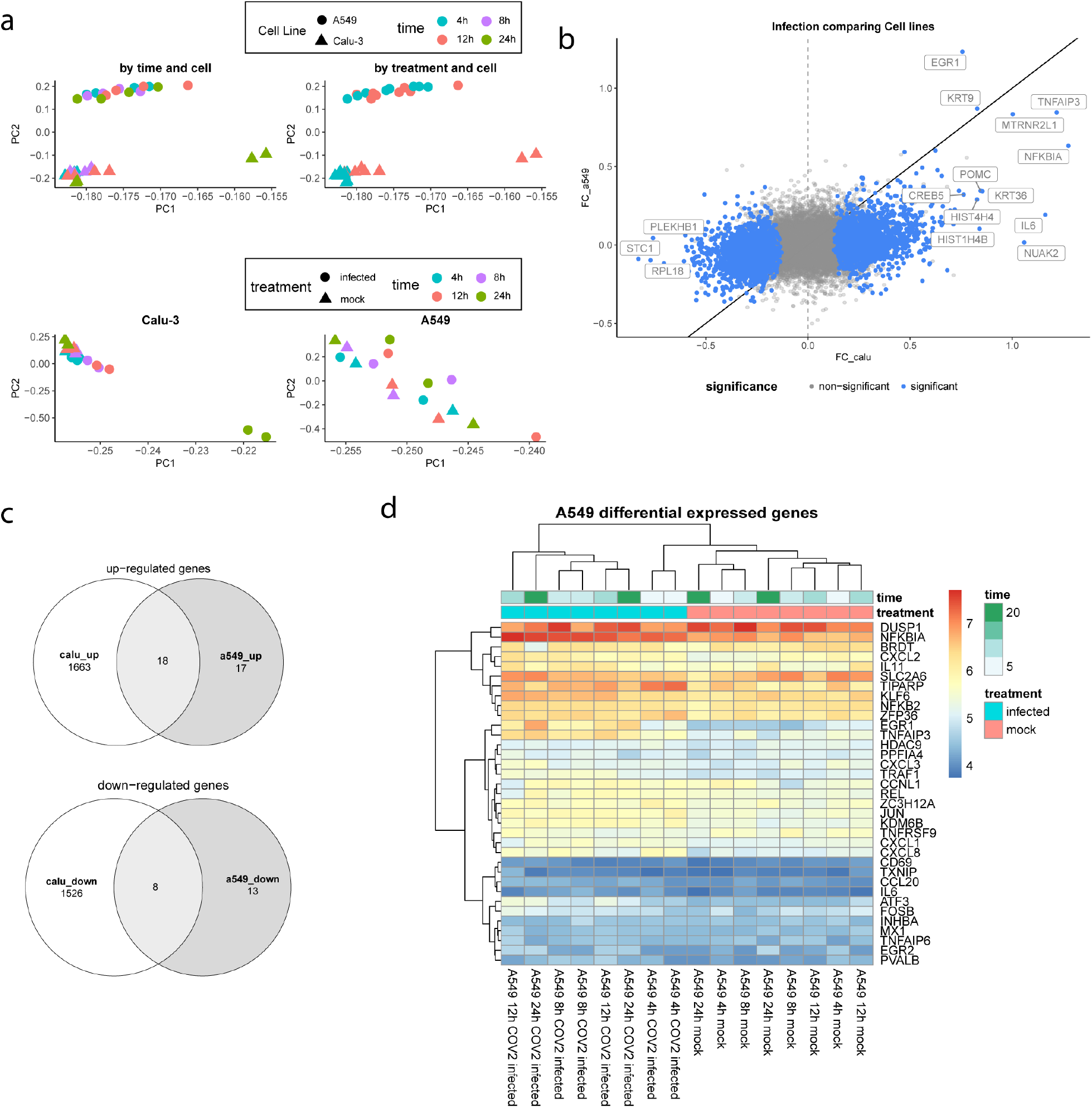
**a,** Principal component analysis on microarray data set. Calu-3 cells show separation between infection and mock but the A549-ACE2 cells have less separation due to lower levels of infection. **b,** Plot comparing enriched genes between the two cell lines. **c,** Venn diagram of significantly (FDR < 10 %) upregulated (top) or downregulated (bottom) genes comparing Calu-3 and A549-ACE2 cells. **d,** Heatmap of enriched genes in A549-ACE2 cells.

**Extended Data Fig. 2.**
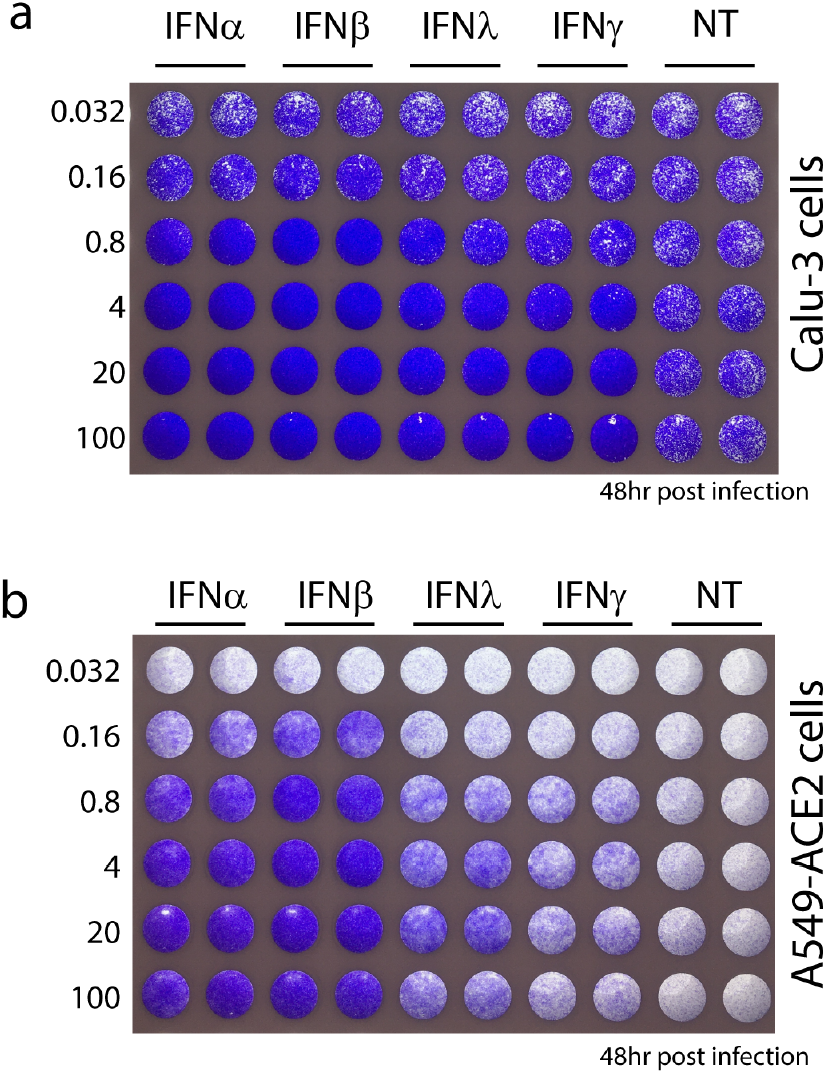
**a-b,** Calu-3 or A549-ACE2 cells were treated with increasing concentrations of given IFNs; highest concentration corresponds to 100X the EC50 value (determined for ISG activation by qPCR) for each IFN. Six hours post treatment cells were infected with SARS-CoV-2 (MOI=5) and 48 h thereafter, cells were fixed in 6% formaldehyde followed by staining with crystal violet.

**Extended Data Fig. 3.**
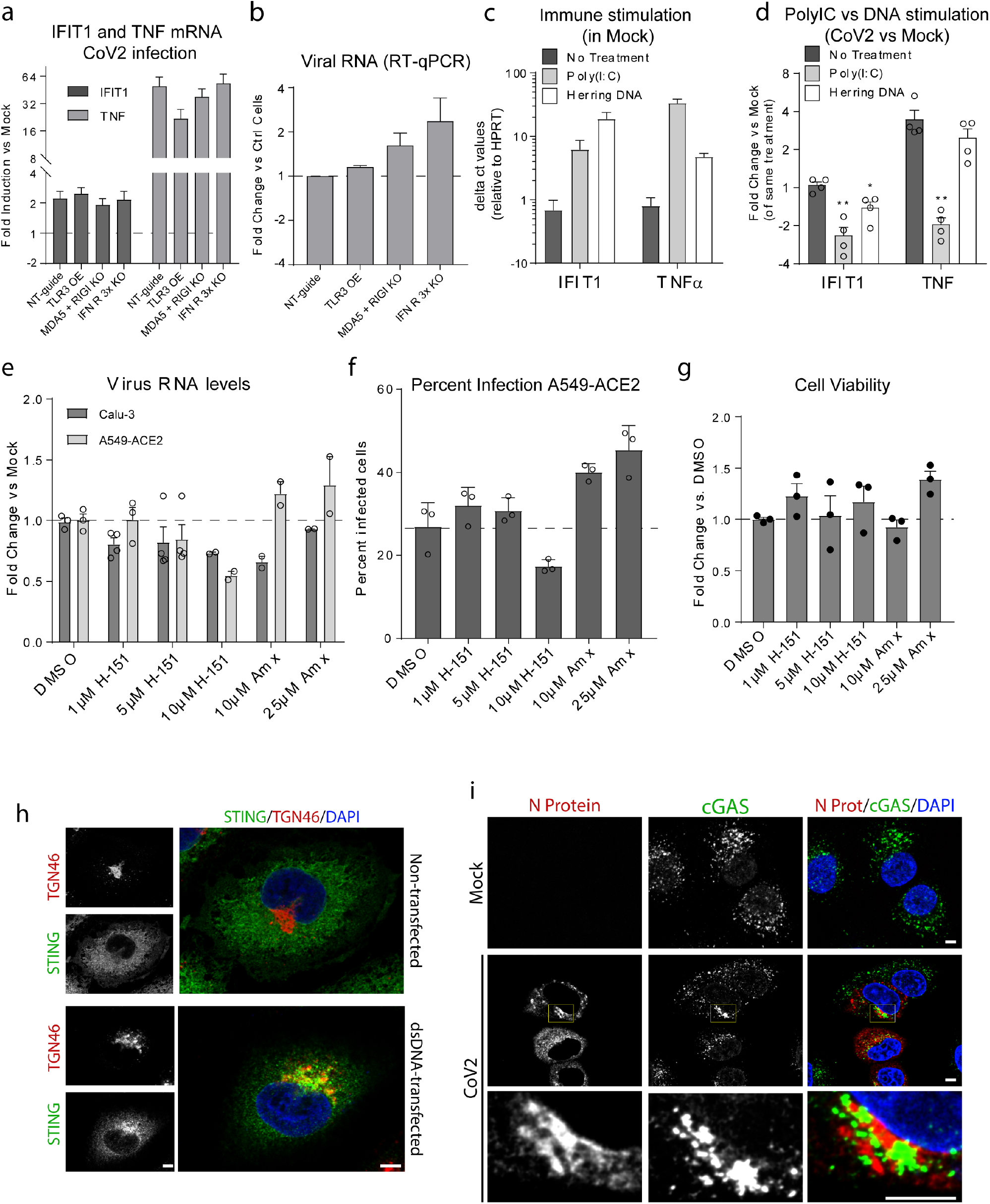
**a-b,** The indicated A549-ACE2 knockout cell lines were infected with SARS-CoV-2 and, 16 h after infection, total RNA was isolated and the levels of IFIT1 and TNF mRNA (a) as well as intracellular viral RNA (b) were determined by RT-qPCR. The graphs show the means and SEMs compared to HPRT mRNA levels for 3 independent experiments. **c-d,** Cells were infected with SARS-CoV-2 or mock-infected for 6 h, then transfected with Poly(I:C) or herring DNA and 4 h thereafter, total RNA was isolated and the mRNA levels of given immune genes were determined by RT-qPCR. **c,** The graphs show the mean mRNA levels of IFIT1 or TNF, corrected for HPRT, for 4 independent experiments for mock-infected cells. **d,** Analogous to panel c, but for SARS-CoV-2 infected cells. Values were corrected for mock of the same treatment for 3 independent experiments. **e-g,** Cells were infected with SARS-CoV-2 and, 1 h later, cells were treated with the given drugs or DMSO only. **e,** total RNA was isolated and the viral RNA levels were determined by RT-qPCR. The graph shows the mean and SEM for 3 independent experiments corrected for HPRT. **f**, Cells were fixed and stained with antibodies specific for dsRNA. Graph shows the mean percent of infected cells from 3 different experiments for each condition. **g,** Graph shows the average fold change and SEM for the number of cells for each drug treatment compared to the DMSO treated cells. **h,** Cells either un-transfected or transfected with herring DNA for 6 h were fixed, stained with the indicated antibodies and examined by confocal microscopy. Scale bar, 5 μm. **i,** A549-ACE2 cells were infected with SARS-CoV-2 for 16 h followed by fixing and staining with the indicated antibodies. Cells were examined by confocal microscopy. Scale bar, 10 μm in the overview and 5 μm for zoom image. Zoomed area is indicated with a rectangle in the middle right panel.

